# A Bayesian Precision Medicine Framework for Calibrating Individualized Therapeutic Indices in Cancer

**DOI:** 10.1101/2021.08.09.455722

**Authors:** Abhisek Saha, Min Jin Ha, Satwik Acharyya, Veerabhadran Baladandayuthapani

## Abstract

The development and clinical implementation of evidence-based precision medicine strategies has become a realistic possibility, primarily due to the rapid accumulation of large-scale genomics and pharmacological data from diverse model systems: patients, cell-lines and drug perturbation studies. We introduce a novel Bayesian modeling framework called the *i*ndividualized the*R*apeutic inde*x* (iR_*x*_) model to integrate high-throughput pharmacogenomic data across model systems. Our iR_*x*_ model achieves three main goals: first, it exploits the conserved biology between patients and cell-lines to calibrate therapeutic response of drugs in patients; second, it finds optimal cell line avatars as proxies for patient(s); and finally, it identifies key genomic drivers explaining cell line-patient similarities. This is achieved through a semi-supervised learning approach, that conflates (unsupervised) sparse latent factor models with (supervised) penalized regression techniques. We propose a unified and tractable Bayesian model for estimation, and inference is conducted via efficient posterior sampling schemes. We illustrate and validate our approach using two existing clinical trial datasets in multiple myeloma and breast cancer studies. We show that our iR_*x*_ model improves prediction accuracy compared to naive alternative approaches, and it consistently outperforms existing methods in literature in both in multiple simulation scenarios as well as real clinical examples.

## 1. Introduction

### 1.1. Scientific background, goals and challenges

Individualized elicitation of therapeutic clinical responses to targeted drug therapies for patients is at the heart of modern precision or personalized medicine endeavors across many disease domains, especially cancer. It is now well-established that patients with similar pathological and diagnostic phenotypes often show varied responses to the same treatment (Ely, 2009; Arjmand et al., 2017). The downstream effects include the risk of adverse drug reactions in patients which in turn makes the therapies costlier (Gray et al., 2018). Thus, for safer and more efficient treatment decisions, understanding patient heterogeneity using the underlying molecular and genetic architecture can aid in prescribing the right treatment to right patients cohort(s) in a precision medicine paradigm (Jiang and Wang, 2010; Kumar-Sinha and Chinnaiyan, 2018). In this context, cancer pharmacogenomics for personalized therapy have been quite successful in developing more accurate and customized diagnostic and prognostic biomarkers in research settings, albeit having limited access in clinical settings (McCarthy, McLeod and Ginsburg, 2013; Farnaes et al., 2018). This motivates deeper investigation of pre-clinical laboratory-based pharmacogenomic models in conjunction with patient’s genomic information to match the right treatment (or combination of treatments) to a future patient, based on their likelihood of eliciting clinical responses.

This is realistically possible, in part, due to the rapid accumulation and availability of large-scale genomic and pharmacological datasets. These include large-scale multi-platform genomics (multi-omics) profiles derived from multiple cancers (e.g. The Cancer Genome Atlas (TCGA, Weinstein et al. (2013)), International Cancer Genome Consortium (ICGC, Zhang et al. (2011)); multi-omics profiles derived from (experimental) model systems in cancer (e.g. Cancer Cell Line Encyclopedia (CCLE, Barretina et al. (2012)), MD Anderson Cell Lines Project (MCLP, Li et al. (2017)), Patient-Derived Tumor Xenograft Encyclopedia (PDXE, Gao et al. (2015)) and large-scale drug perturbation studies in model systems of cancer characterizing their responses to chemical and genetic perturbations (e.g. Library of Integrated Cellular Signatures/ Connectivity Map (LINCS/CMAP, Subramanian et al. (2017)), Dependency Map (DEPMAP, Tsherniak et al. (2017)), Genomics of Drug Sensitivity in Cancer (GDSC, Yang et al. (2012))). This enables systematic analytical approaches to integrate genomic profiles of patients and model systems to uncover novel drug targets, biomarkers, and predictive models in cancer.

Until recently, analytical approaches linking patient and model system genomics data have received less attention from computational researchers, and most previous approaches were developed on patients or model systems alone. For example, high-throughput genomic study of immortalized cancer cell line models and patient-derived xenografts (PDX) models have been successfully used for drug screening, often by identifying statistically significant or predictive biomarkers (Williams and McDermott, 2017; Byrne et al., 2017). In the context of predicting drug response, pharmacogenomic models are often developed on immortalized cancer cell lines – cell cultures undergoing continuous tumor proliferation – as therapeutic biomarker primarily for two reasons. First, large publicly accessible databases (as alluded to above), such as CCLE and, GDSC, allow access to high-throughput drug response across cancer cell lines for many drugs, along with various features of cell lines, including somatic mutations of cancer genes, genome-wide copy number aberrations, and gene expression. The drug response is often measured using IC50 (a measure of potency that indicates how much of a particular drug is needed to inhibit a given biological process by half) or AUC (The area under the plasma drug concentration-time curve (AUC) that reflects the actual body exposure to the drug after administration of a dose). In contrast, clinical trial data or other patient-derived models (such as PDX), are often very expensive and, thus, not available for variety of drugs. Second, many recent studies stress the similarity between cell lines and patient data in terms of common predictive “-omic” features/bio-markers (Domcke et al., 2013; Sinha et al., 2017). Most of these models are trained on cell lines and tested (independently) on patient data from clinical trials (Geeleher, Cox and Huang, 2014; Falgreen et al., 2015).

In many instances, however, although genomic data for patients are available in the beginning of or during clinical trials, they often remain unused for training such models. Thus, these models suffer from a fundamental drawback that training samples and test samples come from two different sources. This invariably leads to bias and out-of-sample prediction on patients that can be severely influenced, if patients’ genomic measurements significantly differ from that of the cell lines. This is possible when cell line samples evolve quite differently from patients at a molecular level, in absence of the stromal eco-system surrounding tumors (Wang et al., 2006; Brodaczewska et al., 2016). Such source-specific variations would impact any analysis pipeline that treats the samples alike. Hence, quantification and delineation of both the source-specific and conserved variation between patients and cell lines is critical. Many studies have shown that some key oncogenic programming is conserved across models, as well as cancer states (Piñeiro-Yáñez et al., 2018; Califano and Alvarez, 2017). Thus, exploiting this conserved biology between patients and cell lines is key to understanding therapeutic response for patients based on drug perturbations in cell-lines. This motivates us to build an integrative methodology for patients, cell-lines and drugs, that serves three key purposes: a) to use patient-cell line similarities to predict therapeutic response, b) to find “optimal” cell line avatars as proxies for patients (or groups of patients), c) to find key genomic drivers (e.g., pathways, sets of genes) explaining patient-cell line similarities.

### 1.2. Overview of our model

In this article, we propose a novel Bayesian semi-supervised framework, called the <u>i</u>ndividualized the<u>R</u>apeutic inde<u>x</u> (iR_*x*_) model, that achieves the above goals. A conceptual schematic of the iR_*x*_ model is shown in Figure 1 (explicit details are presented in Section 2.1). Briefly, we build a joint model on the entire {patient, cell-line, drug} data triplet by jointly specifying two sub-models: the *Drug-cell line* model (Model A) that relates drug responses to cell lines’ genomic data, and the *Cell line-patient* model (Model B) that explicitly models the relationship between genomic profiles from the two model systems. The Drug-cell line model serves as our (labeled) training model, which follows a penalized regression model. The Cell line-patient model captures the underlying genomic similarity using sparse low-dimensional latent factor models, explicitly parametrized to model and quantify source specific and shared variations separately. These latent factors essentially capture the underlying conserved biological process across the two model systems. The sub-models inherit nice interpretive and computational properties. In essence, the iR_*x*_ model conflates (unsupervised) latent factor models with (supervised) penalized regression techniques, which engenders a semi-supervised method to calibrate (missing) responses for patients, called iR_*x*_ scores (Model C), borrows information from both the Drug-cell line and Cell line-patient models.

**Fig 1:**
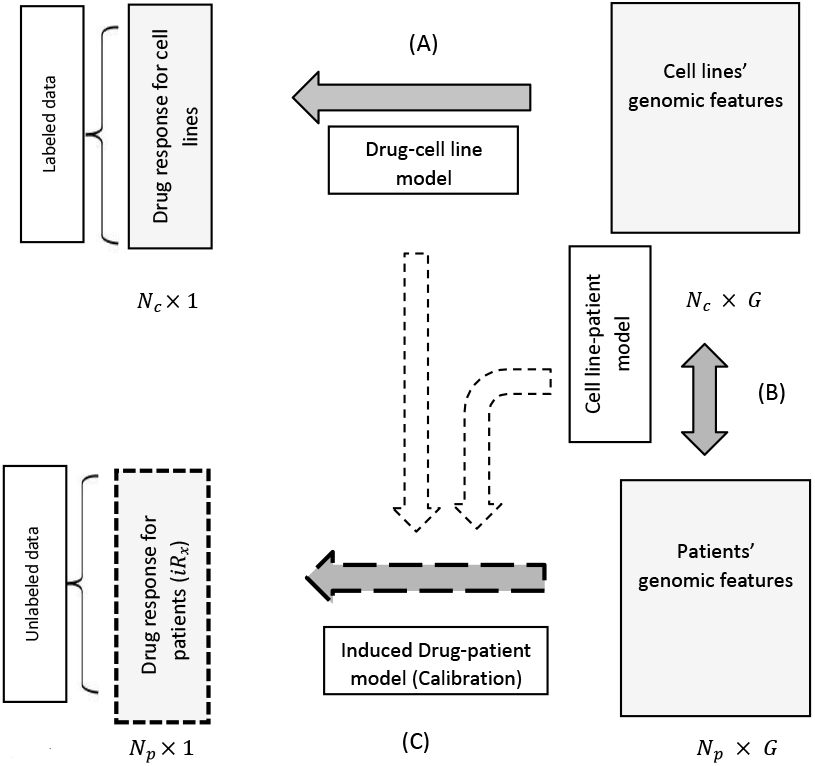
Conceptual framework of the iR_*x*_ model. The joint model is specified by two sub-models, namely the *Drug-cell line* model (A) specifying the relationship between drug response (*N*_*c*_ × 1 vector) and the cell lines’ genomic features (*N*_*c*_ × *G* matrix), and the *Cell line-patient* model (B) specifying the relationship between genomic features from two sources. These are then integrated to calculate the iR_*x*_ score (*N*_*p*_ × 1) using the induced *Drug-patient* model (C). *N*_*p*_, *N*_*c*_ and *G* represent samples sizes of patients, cell lines, and number of matched genes, respectively,

We illustrate and validate our approach, retrospectively, using two existing clinical trial datasets in cancer: a bortezomib study in multiple myeloma patients (Mulligan et al., 2007), and a docetaxel study on breast cancer patients (Chang et al., 2003) – wherein the patient-specific drug responses are available, that are blinded for modeling and only used for model validation. We show that our iR_*x*_ model improves prediction accuracy compared to naive alternative approaches and consistently outperforms existing methods in literature, in both the real data examples and in multiple simulation scenarios. In addition, we identify key cell-line “avatars” of patients (Figure 3) and key clinically relevant signaling pathways that drive the conserved biology between patients and cell-lines (Figure 5).

Sections 2.1 and 2.2 introduce the fundamental inference problem, as well as the statistical model and its interpretations. Section 2.3 introduces the calibrated estimands: iR_*x*_, and, its properties. Section 3.1 to 3.3 delineate the Bayesian estimation process, along with priors and sparsity assumptions and Markov chain Monte Carlo (MCMC) computations. Section 3.4 introduces the clustering problem, along with theoretical justifications. Section 3.5 elaborates on gene and pathway selection. Section 4 illustrates two sets of simulation studies, and Section 5 summarizes our real clinical trial data analyses. Finally, Section 6 provides our conclusions and discussion.

## 2. The iR_*x*_ Model

### Data formats

Following the conceptual schematic in Figure 1 for the iR_*x*_ model, the specific data format considered is as follows. For a given drug *D*, assayed on *N*_*c*_ cell lines, let 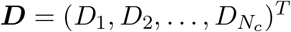 be the *N*_*c*_ dimensional vector containing the drug response measurements (such as IC50 and AUC). We denote the genomic profiles for the cell lines as a *G* × *N*_*c*_ matrix 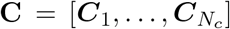, where *C*_*j*_ is a *G*-dimensional vector having genomic variables (e.g., genes, proteins, mutations) for *g* = 1, … , *G* of the *j*^*th*^ cell line. Our primary objective is to calibrate corresponding drug response on a patient cohort of size *N*_*p*_, using the measured genomic profiles of the same set of genomic variables, *V* = {1, … , *G*} on patients. The corresponding measured genomic profiles on *G* genes in *V* for patients are denoted by *G*× *N*_*p*_ matrix 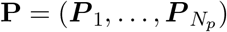, where *P*_*j*_ is *G*-dimensional vector having measurements for *G* genomic variables on the *j*^*th*^ patient. Succinctly, the (observed) data available can be summarized as a tuple: (***D*, *C*, P**). In this paper, C and P include transcriptomics (mRNA expression), but other’omic platforms (proteomics, epigenomics, mutations) can easily be used as well as across multiple drugs (see Section 6).

### 2.1. The basic inference problem: labeled and unlabeled data

We cast the basic inferential problem in a semi-supervised learning framework (Chapelle, Scholkopf and Zien, 2006), wherein we have two sets of data: labeled and unlabeled:

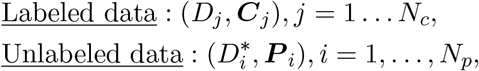

where (*) represents the unobserved labels (patients’ drug response). Thus, the basic task of calibration of drug responses for patients reduces to estimating 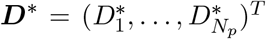 based on the available data (***D*, C, P**) (i.e., to develop a predictive model to calibrate the unobserved drug responses (labels)).

Suppose **Θ** represents the union of all underlying parameters of interest, which are explicitly elaborated in the next Section. The labeled data, {(*D*_*j*_, ***C***_*j*_), *j* = 1 … *N*_*c*_, are assumed *i.i.d* samples from the underlying distribution, 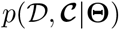, where 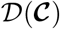 is a generic random variable (vector) for *D*_*j*_ (*C*_*j*_), *j* = 1, … , *N*_*c*_. Analogously, 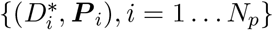 can be treated as *i.i.d* samples from the underlying distribution, 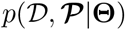. Hence, the inference for a drug response on a given patient with genomic profile 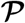 will be based on 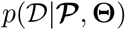 – which is our target distribution of interest, for inference and prediction.

### Distributional Assumptions

For a generic patient with genomic features 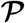, the target distribution can be expressed as,

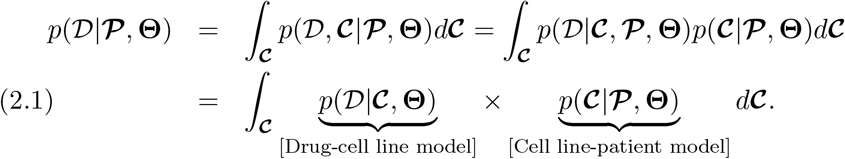

In essence, 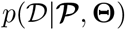 can be learned by the conflation of two sub-models: a *Drug-cell line model*, 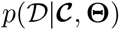, and a *Cell line-patient model*, 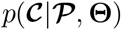. Note, in the second equation we assume 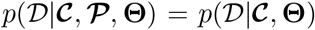, which implies, given the features of both cell lines and patients, drug sensitivity scores depend on only the features of cell lines and the commonality shared between cell lines and patients through **Θ**. Thus, **Θ**, in general, allows further sharing of information between cell line (**C**) and patient (**P**) data, through 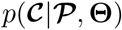 model, and thereby posterior predictive density, 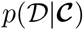 would depend (latently) on **P**. We believe the assumption is quite realistic as 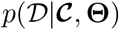 only ignores any dependence due to 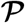, which is other-wise accounted in patient-specific idiosyncratic errors in 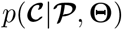 model. This is also biologically justified due to: a) the nature of data collection, since cell lines are the only ones labeled with drug information; and b) there is considerable literature support in pre-clinical cancer studies, that cell line systems could show significant differences in their growth, compared to patient tumors, primarily due to absence of stromal cells surrounding the tumor (see Wang et al. (2006); Brodaczewska et al. (2016) and references there-in). Thus, note that the drug-cell line model involves only the labeled data, while the cell line patient model involves the unlabeled data. Following the conceptual schema in Figure 1, where the top Section represents labeled data (Drug-cell line model) and bottom right part is unlabeled data (patients’ features), the solid yellow Section represents the observed drug response (labels), whereas the dotted yellow Section is unobserved drug response that needs to be estimated in the iR_*x*_. Figure 1 also shows two sub-models through blue solid arrows ((A) and (B)) and calibration method through the dotted arrow ((C)), that in turn depends on two sub-models. We will show later that pooling information across labeled and unlabeled data (by using Cell line-patient model) improves calibration accuracy compared to the more common alternative that uses only Drug-cell line model, both in simulated and real data applications (Sections 4 and 5). In the ensuing Sections, we specify each of the sub-models, along with the semi-supervised estimation procedure specified in (2.1).

### 2.2. Model specifications

#### Drug-cell line model

For the drug cell-line model 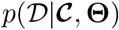, we propose a regression formulation, where the drug responses are regressed upon cell lines’ genomic features. Given a continuous drug-response, the form can be expressed as,

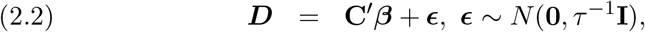

where *β* = (*β*_1_, *β*_2_, … , *β*_*G*_)^*T*^ represents *G* × 1 vector of regression coefficients and 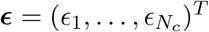 is a *N*_*c*_× 1 residual vector. In this linear model setting, *β*_*g*_ represents the effect of the *g*^*th*^ gene for *g* = 1 … *G*.

There are two main factors that govern the choice of the regression formulation for the drug-cell line model. First, it allows for building an explicit predictive model on the training data from the genes obtained on the cell lines. Second, it thus allows drug sensitivity to depend on patients by borrowing strength from the cell line-patient model (discussed next) and consequently facilitating explicit calibration of iR_*x*_ scores (Section 2.3). Furthermore, from a computational standpoint, since we are in a high-dimensional setting (*G* ≫ *N*_*c*_), we use a penalized estimation approach using ridge regression (Hoerl and Kennard, 1970), which penalizes ∥*β*∥, where ∥.∥ represents an *L*_2_ norm. This penalization on *β* induces shrinkage estimators, as well as balances predictive power (Bøvelstad et al., 2007; Stransky et al., 2015). We defer the details of this estimation procedure until Section 3, after we complete the full model specification. We note here that a (Gaussian) linear regression model suffices for our case study, but any non-linear functional forms could be used (see Section 6).

#### Cell line-patient model

Instead of specifying the conditional model of cell lines’ genomic measurements given patients’ profiles, 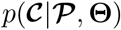, we first specify a joint model for *p*(**C, P**|**Θ**), where **C, P** are the data matrices for the i.i.d samples collected corresponding to random vectors, 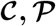, of a single sample respectively, that would in-turn induce conditional model required for iR_*x*_ computation (see Table 1 of Appendix A to know more about various notations used). This allows for modeling the common dependence structure within genomic features, while accounting for cell line-specific and patient-specific variations. To this end, we propose the following latent factor model

**Table 1.**
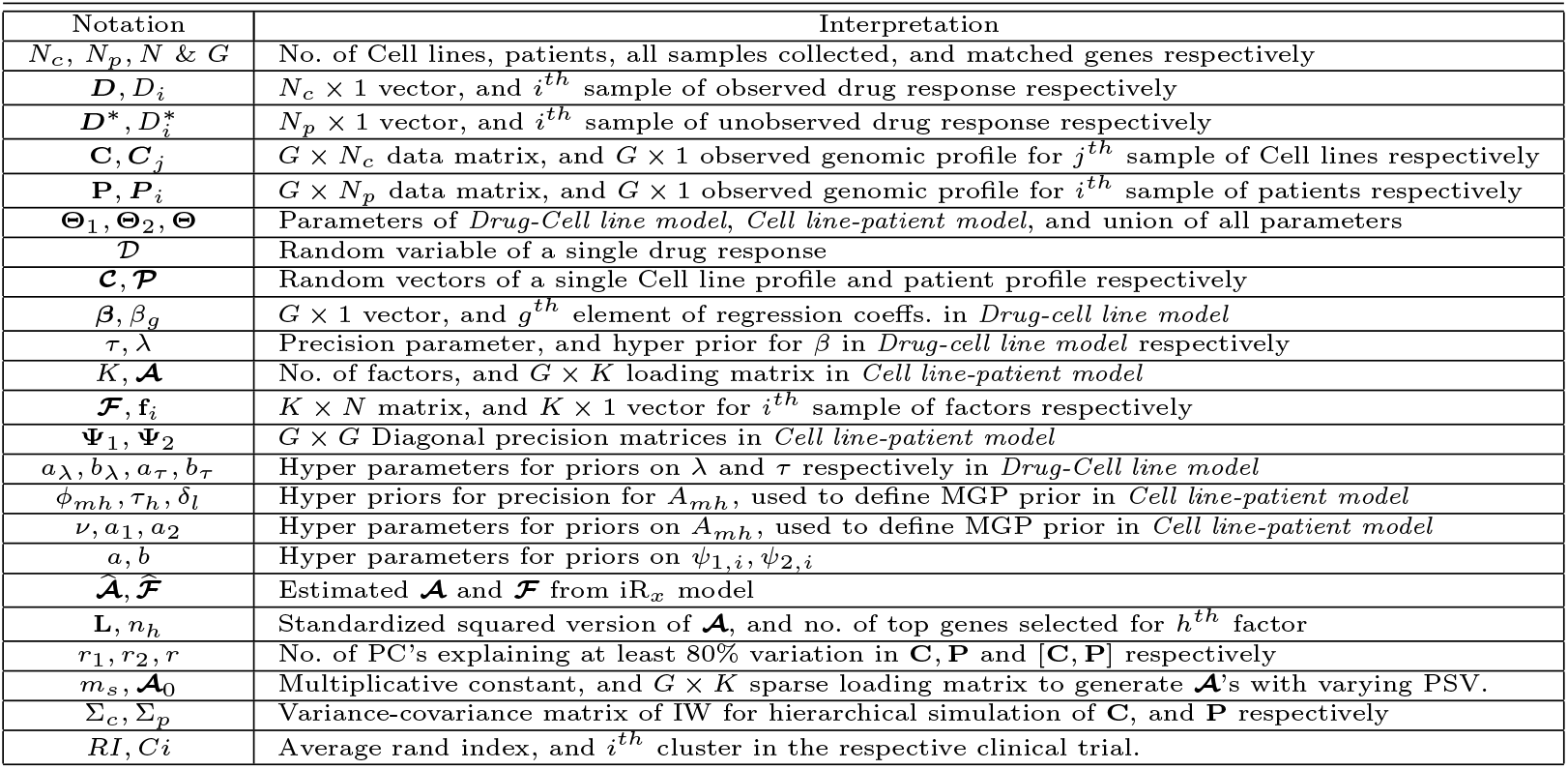
Notations used in the manuscript along with their interpretations

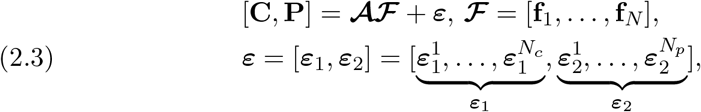

where [**C, P**] is a horizontally pooled gene expression matrix of dimension *G* × *N, N* = *N*_*c*_ + *N*_*p*_, total sample size. For *K* ≪ *G*, 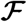 is the *K* × *N* matrix of f_*i*_’s for *i* = 1, … , *N*, where f_*i*_ is a vector of *K* factors for *i*^*th*^ sample, and 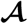 is the *G* × *K* loading matrix that is shared across patients and cell lines, where the row vector, 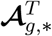is the *K* × 1 vector of loadings corresponding to the factors for gene *g* in *V*. Following a standard factor model, we assume that the vectors of factors, f_*i*_’s, are identically and independently distributed as *N* (0, **I**_*K*_) for *i* = 1, … , *N*. The *G* × 1 error vectors for cell lines and patients, 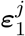 for *j* = 1, … , *N*_*c*_ and 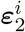 for *i* = 1, … , *N*_*p*_, are assumed to follow 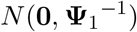 and 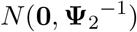, respectively, where **Ψ**_1_ and **Ψ**_2_ are unknown diagonal precision matrices that represent idiosyncratic noises. We further assume that *ε* and 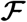 are mutually independent.

#### Dependence within genes via latent structure

We induce the dependence structure of genes by the factor model. For example, for the *j*^*th*^ cell line sample, we take measurements of two genes, *g* and *g*′, denoted by *C*_*g,j*_ and *C*_*g′,j*_. From the model in (2.3), we represent them by linear models of common latent factors: 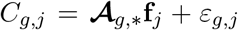 and 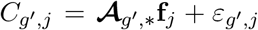, where 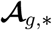 and *ε*_*g,j*_ are the corresponding elements for *g*^*th*^ gene. Since f_*j*_ ~ *N* (**0, I**_*K*_), the induced covariance between the two genes is 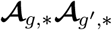, that is common for cell lines and patients. Thus, *G*-dimensional genomic data for a patient and a cell line are expressed as 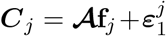 and 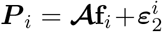 with covariance structures, 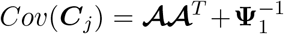 and 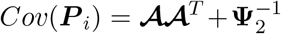. Thus by sharing the factor loading matrix 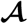, we posit a common (genomically driven) dependence structure between cell lines and patients, whereas the distinct residual variances of genes between cell lines and patients, 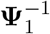 and 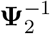, account for the variation in genes that are not explained by this common, shared variance structure. In this article, we assume that the loading matrix 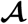 is sparse, which also makes it identifiable (Lucas et al., 2006).

### 2.3. Individualized theRapeutic indeX (iR_x_) calibration

First, to compute 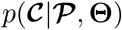, the induced cell line-patient conditional model, we are required to specify the joint distribution between 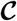 and 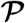. Following (2.3), 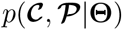 can be expressed as,

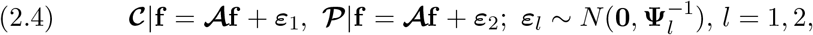

where f, the common *K*-dimensional vector of factors, follows *N* (**0, I**_*K*_). *ε*_1_, *ε*_2_ and f are independently distributed (as above). Using (2.3) and (2.4), one can obtain our target distribution of interest, containing information on drug response for patients 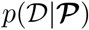, as

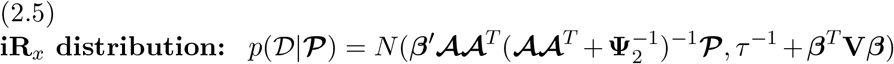

where V is 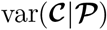 (see Section S1 of the Supplementary Materials for derivation). Since we have characterized the entire distribution of iR_*x*_, there are various ways to extract summary measures of drug response for a given patient from this distribution (e.g., any moments or quantiles). A natural way would be to consider 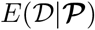, which is what we term as an *i*ndividualized the*R*apeutic inde*x*, or iR_*x*_ score, and can be construed as the best average drug response for a given drug 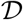. This leads to the following proposition that defines the iR_*x*_ score of a drug for a patient with a genomic profile 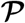 as:

#### Proposition 1.

*Under the proposed model in (2.2), (2.3) and (2.4), for a patient with genomic profile* 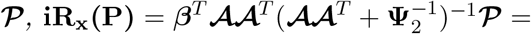 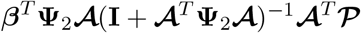

Proof of Proposition 1 is given in Section S1 of the Supplementary Materials. In practice, for the *i*^*th*^ patient sample with genomic measurements *P*_*i*_,

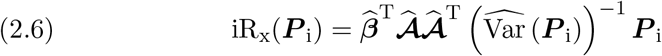

where 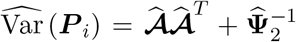, and 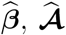, and 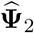 are the corresponding estimates.

### Naive Index (NI)

As an alternative to iR_*x*_, we define a naive index, **NI**, solely based on the Drug-cell line model in (2.2), as 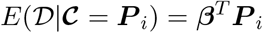. Thus, NI is given by,

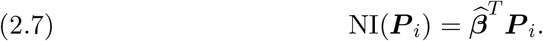

To compare these scores, we consider the mean squared prediction error (MSPE) for a given patient *P*_*i*_. Since our target is to estimate drug response 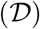, for an estimator 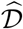, 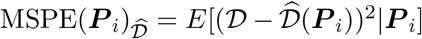. With the two choices for 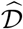, namely iR_*x*_ and NI, we make the following claim.

### Claim

*Under the assumption of shared genomic variation across cell lines and patients as in (4)*, 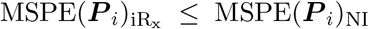. *In addition, gain in accuracy is given by* 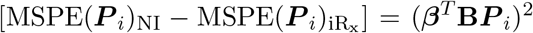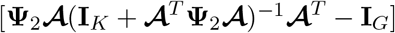.

Proof of Claim is provided in Section S1 of the Supplementary Materials. The claim ensures that the (prediction) accuracy of iR_*x*_, as measured by MSPE, always improves upon that of NI, when there exists a shared genomic dependence structure between cell lines and patients, where-in the gain in accuracy can be given by the term (*β*^*T*^ **B*P***_*i*_)^2^.

### Standardized iR_*x*_ score and dichotomized iR_*x*_ classifier

For iR_*x*_ scores to be comparable across various patient groups (e.g., across various subtypes or stages of cancer), we can work with a standardized iR_*x*_ score, or 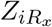 score, where *Z*_*i*_ = [iR_x_(*P*_i_) − *μ*]/*σ*, where *μ* = (1/*N*_*p*_)Σ iR_x_(*P*_i_), and *σ* is the empirical standard deviation across the patient-group under consideration. This puts the iR_*x*_ scores on a common scale for comparative analyses (e.g., evaluating drug responses across multiple patient groups) for a clinical investigator. On the other hand, there might be instances where a dichotomous score will be more interpretable for classification (e.g., if a specific patient responded or not to an explicit therapy or drug). To this end, one can use a dichotomized iR_*x*_ classifier, based on the iR_*x*_ score, to distinguish between responders and non-responders. We propose a natural classifier based on a *cσ* bound, which can be defined as: *Z*_*i*_ > *cσ* ⇒ non-responder, *Z*_*i*_ < −*cσ* ⇒ responder, and |*Z*_*i*_| ≤ *cσ* ⇒ no effect. The choice of *c* can be based on the specific patient population being modeled and may depend on patient-specific characteristics (e.g., cancer type, stage of the cancer).

## 3. iR_*x*_ model estimation and inference

While estimation and inference of iR_*x*_ model, as specified in Section 2.2 and 2.3, can be done via frequentist (e.g., optimization, expectation-maximization) or Bayesian techniques, we propose a unified fully Bayesian formulation, since it allows us to incorporate a rich and flexible class of prior structures for inducing sparsity and regularization. Let **Θ**_1_ and **Θ**_2_ be {*β, τ*}, the set of parameters of the drug-cell line model, and 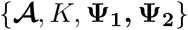, the set of parameters for the cell line-patient model, respectively. Further define the set of all parameters, **Θ** = **Θ**_1_ ∪ Θ_2_. Under the assumption that parameters of two sub-models are independent to each other (i.e., *p*(**Θ**)= *p*(**Θ**_1_)*p*(**Θ**_2_)), the posterior of **Θ** is given as follows:

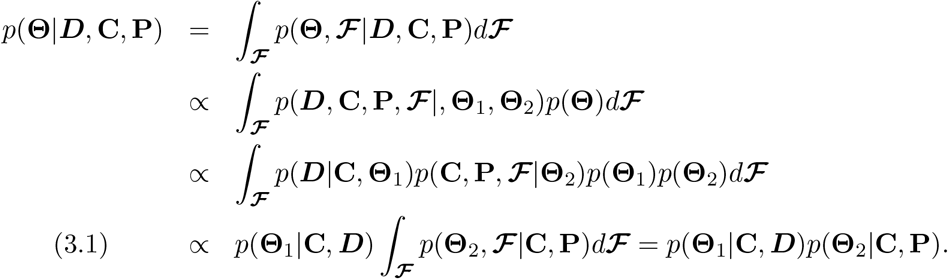

If we specify the priors of **Θ**_1_ and **Θ**_2_ independently, the posterior of **Θ** factors into posteriors of **Θ**_1_ and **Θ**_2_. Thus, we next specify priors independently for each model and implement a Markov Chain Monte Carlo (MCMC) based sampler for estimation and inference.

### 3.1. Drug-cell line model

For estimation of (2.2), a Bayesian ridge model is considered, priors for which can be given as follows.

*p*(**Θ**_1_) = *p*({*β, τ*}) = *p*(*β|τ*)*p*(*τ*), *p*(*β|τ*) ~ *N*(**0**, [*λτ*]^−1^***I***), 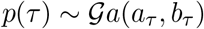, where *λ* is a hyper parameter for *β*, and *a*_*τ*_ and *b*_*τ*_ are hyper-parameters for *τ* (Gelman et al., 1995). This method essentially applies a shrinkage prior that puts high precision around 0 on the regression coefficients for large *λ*. Thus, a high *λ* would shrink the estimates more. We let the data determine shrinkage by assuming *λ* follows 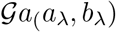, where *a*_λ_, *b*_λ_ are chosen in such a way that gamma distribution remains weakly informative, yet proper. Choices for *a*_*τ*_, *b*_*τ*_ are made the same as *a*_λ_, *b*_λ_.

### 3.2. Cell line-patient model

Cell line-patient model in (2.3) gives rise to a standard sparse factor model when **Ψ**_1_ = **Ψ**_2_. Due to high dimensional nature of loading matrix 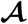, estimation requires methodologies that adaptively induce sparsity and automatically remove redundant parameters not required to characterize the data. In this regard, we use the multiplicative gamma process (MGP) (Bhattacharya and Dunson, 2011) shrinkage prior, as it leverages an infinite representation of 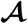 to start with and induces increasing shrinkage on loadings as the number of factors increases, thereby choosing a parsimonious model with a much smaller effective number of significant factors (say, *K*). The MGP prior is defined on 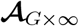 and *K* is determined by eventually keeping left most significant columns of 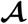. Math-ematically, the MGP prior can be given as follows: for 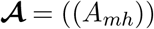,

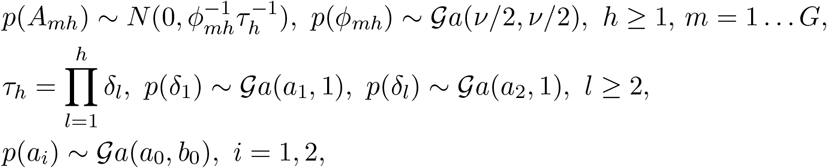

where *δ*_*l*_, *l* = 1 … ∞ are i.i.d; *φ*_*mh*_ is the local shrinkage parameter for *h*^*th*^ column and *m*^*th*^ gene and *τ*_*h*_ is the global shrinkage parameter for *h*^*th*^ column of the loading matrix. *τ*_*h*_ is stochastically increasing for *a*_2_ > 1. As a result, shrinkage on the column increases exponentially with the column index, leading to first few factors from the left, having significant loadings only. Diagonal elements of precision matrices, **Ψ**_1_ and **Ψ**_2_, are assumed to have independent gamma priors, 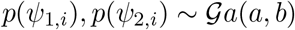, *i* = 1 … *G*, where *a, b* are chosen, as is done in the case of *p*(*λ*). This completes the specification of the prior for 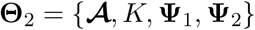.

### 3.3. MCMC computations

For estimation and inference, we first compute full posterior likelihood, and subsequently full conditionals for gibbs sampling steps. As shown in (3.1), estimation of the set of all parameters, 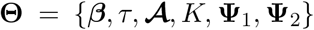 can be done efficiently, since drawing samples from *p*(**Θ**|***C, D***) is equivalent to drawing samples of *p*(**Θ**_1_ = {*β, τ*}|**C, *D***), independently to drawing from 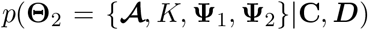 due to fac-torization. As a result, MCMC samplers for two posteriors, *p*(**Θ**_1_|**C, *D***) and *p*(**Θ**_2_|**C, *D***), are run in parallel. In each case, we have used the posterior means of the samples to estimate the corresponding parameters. The full posterior for *p*(**Θ**_1_|**C, *D***) is computed using a collapsed Gibbs sampler, while for posterior computation of *p*(**Θ**_2_|**C, P**) we employ an adaptive Gibbs sampler, proposed in Bhattacharya and Dunson (2011), after accommodating for an additional precision matrix in our setting. Full details on the joint posterior, the full conditionals, and the implementation strategy are given in Section S2 of the Supplementary Materials. As for estimation of *K*, we note that posterior mean and posterior mode are the same as the value of *K* stabilizes to a single number very fast after a few initial iterations (see Figure S2 of the Supplementary Materials). We then compute both iR_*x*_ and NI scores using (2.6) and (2.7). In addition to storing parameter estimates, we also store the posterior means of factors, on which K-means method is applied with the estimated number of factors, *K*, to obtain clusters based on genomic similarity (discussed next).

### 3.4. Patient Avatars based on genomic similarity

The integrative modeling across cell lines and patients (via the cell line-patient model) provides information on similarity between them due to underlying genomic or transcriptomic features. Since there is some preservation of the steady state cancer cell behaviors across model systems (e.g., common oncogenic or transcriptional programs Califano and Alvarez (2017)), we hypothesize patient tumors and cell lines having similar underlying mechanistic features that would show similar therapeutic responses to the same drug. Thus, if cell line(s) are clustered together with patients based on this similarity, they can be studied for therapeutic implications as proxies for the patient subgroup from the same cluster. Such cell line(s) representing a patient sub-group are termed as avatars for the corresponding patient sub-group. To obtain patient avatars, we propose a clustering algorithm based on shared variation due to underlying transcriptomic features from the latent factor model.

Let us denote the expression matrix of estimated samples based on shared variation as 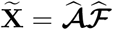, where 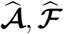 are estimated from the iR_*x*_ model. One can also interpret 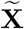 as X adjusted for source-specific variations, which hence would give desired clusters (via *K*-means) of patients and cell lines based on shared variation. The following remark assures us that in order to obtain such clusters, it is enough to run a *K*-means clustering on estimated factors, *K* being the number of factors.

#### Remark

*K-means clustering on estimated factors (posterior mean)*, 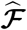(*i.e. on the columns of* 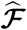) *obtained from (2.3) will approximately produce the same clusters as K-means clustering based on shared variation (on the columns of* 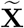), *with K, number of clusters, being the number of factors estimated*.

The justification for the choice of number of clusters and the proof of **Remark** are given in the Section S3 of the Supplementary Materials. The clustering mechanism enjoys several advantages: (1) like other spectral clustering methods, it is computationally fast, since typically *K* is of moderate dimension; (2) it does not require tuning for *K*, since it is automatically chosen by the MGP prior, and (3) avoids computation of 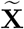 explicitly, which can be expensive. We employ this algorithm to our real data and focus on the mixed clusters, having both cell-lines and patients, to interpret the avatars in terms of drug response phenotypes (see Section 5).

### 3.5. Identification of key genomic drivers of shared variation

In the latent factor model, (2.3), each latent factor represents a potential biological mechanism, which can be used to identify key genomic drivers of the shared variation between patients and cell lines. Specifically, the factor loadings of the *h*^*th*^ column, 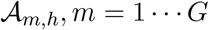, represent each individual gene contributions to the *h*^*th*^ factor. Hence, by evaluating the patterns of factor loadings, we can gain understanding of the biological mechanism common to both cell lines and patients genomic data. Furthermore, the significant loadings help with summarizing genes that act as surrogates for the underlying biological mechanisms. Thus, for every factor, we are interested in working with the top *n*_*h*_(< *G*) significant genes that better represent the factor.

To this end, we need a selection criterion that introduces sparse loadings by thresholding at significant genes and making them comparable across factors. To achieve this, we consider the standardized squared magnitude of loadings,

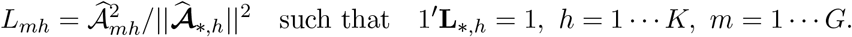

Further, we threshold to the top values that account for 95% of the total. In other words, if *n*_*h*_ top genes are selected for factor *h*, then 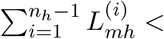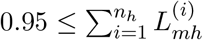, where 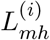 is *i*^*th*^ ordered standardized loading. Thus, for every column *h*, the other genes accounting for 5% contribute to the sparsity of the column. The sparsity increases from the left-most factor to the right-most factor as *n*_K_ ≪ *n*_1_. In practice, this may indicate that some biological phenomena may be easier to detect, wherein few genes play significant roles due to highly sparse columns as we move towards the right. In real data analysis, we used a standard pathway analysis tool to associate the selected genes to potentially clinically relevant pathways (see Section 5).

## 4. Simulation Study

We have considered two types of scenarios for our simulations. In Simulation Study I, we compared the prediction accuracy of our iR_*x*_ method with other competing methods in simulation settings with various degrees of heterogeneity between the cell lines and patients molecular profiling data (Section 4.1). We also investigated the robustness of our clustering procedure to the estimated number of factors. In Simulation Study II, we emulated the Myeloma study (Section 5.1) by generating datasets from the estimated cell line-patient model (Section 4.2). We focused on addressing the identifiability of our cell line-patient model as well as evaluating the prediction accuracy.

The design parameters that are constant across scenarios include: number of genes, *G* = 1000; and the precision parameter in (2.2), *τ* = 0.25; and the regression coefficients, *β*_1_, … , *β*_*G*_ ~ *N* (0, 1) as the estimates from the multiple myeloma study are (approximate) standard normal draws (see Section S4 of the Supplementary Materials). In each setting, we generated 50 replicates of {***D*, C, P**}, and *D*^*^ using (2.2), (2.3), and (2.5), respectively. For estimation, both the (collapsed) Gibbs sampler for (2.2) and the adaptive Gibbs sampler for (2.3) are run in parallel in which the first 5000 iterations were burn-in, and then the chain was thinned by drawing one in every 5^*th*^ iteration thereafter. Posterior means (median for *K*) are used as parameter estimates. The convergence of the MCMC chain was assessed by trace-plots and standard MCMC diagnostics (details in Section S5 of the Supplementary Materials). Since the choices for initialization of parameters and choices of hyper-parameters remain the same across simulation studies and real data applications, these details are given in Section S2 of the Supplementary Materials, along with the corresponding posterior distributions and all codes provided to reproduce the results.

### 4.1. Simulation Study I

In this study, the true models considered for data generation are of two types, namely, a) Scenario 1: Known structure of shared variation, b) Scenario 2: Unknown structure of shared variation. We compare the performance of iR_*x*_ to other competing methods: the NI method, and two principal component (PC) regression based methods (similar in spirit to those Ma et al. (2018)), namely `PC`_ind_, `PC`_joint_ that use a fixed set of principal components. PC regression is performed on ***D*** based on PCs of **C. *D***^*^ is predicted by extending regressions upon PCs of **P**. `PC`_ind_, `PC`_joint_ differ from one another in the manner the top PCs are computed and chosen. For `PC`_ind_, top PC’s of **C** and **P** are computed separately so that we choose *r*_1_ and *r*_2_ of them such that they each explain 80% variation in their data-sets. We propose to work with top *r* = max(*r*_1_*, r*_2_) many PC’s from each data-set. In presence of some shared structure, top PCs of corresponding order can be shown to be strongly correlated (Figure S11, Section S11.1 of the Supplementary Materials). In `PC`_joint_ method, we compute PC’s based on the joint matrix [**C,P**] and work with top *r* many PC’s that explain 80% variation in [**C,P**].

In essence, in Scenario I all three methods except NI assume implicitly, a common structure of shared variation and assesses iR_*x*_’s relative gain in accuracy relative to other methods. Scenario I allows us to study the stability of the number of factor and the clusters estimated in iR_*x*_ model under various cases of heterogeneity considered in shared variation as well as in source specific variations. Scenario II complements the Scenario I, and evaluates the robustness of these methods are in absence of shared structure.

#### 4.1.1. Scenario I: Known structure of shared variation

Since, in general, to estimate the prediction accuracy, true scores, ***D***^*^ or *p*(***D*|P**) needs to be known, an arbitrary dependence structure of *p*(***D*|P**) on **C** may not result in shared variation in a desired manner. To evaluate the claim of gains in prediction accuracy of iR_*x*_ (in Section 2.2.) in presence of high shared variation relative to source-specific total variations, it is worthwhile to consider scenarios where shared variation is varied from a low proportion through a high proportion in relative to average total source specific variations. To achieve this, we first define shared variation by 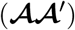 while total source specific variation by 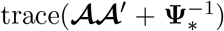. To keep the settings simple and easy to interpret, we control variation in 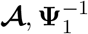 and 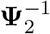 by three parameters, *m, ψ*_1_ and *ψ*_2_ respectively such that 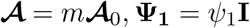, and **Ψ**_2_ = *ψ*_2_**I**, where 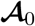 is *G* × *K* sparse matrix given choices of *ψ*_1_ and *ψ*_2_. Note that adding*ψ*−1**I** for *ψ* > 0 to a variance-covariance matrix of dimension *G* is equivalent to increasing the total variation by *Gψ*^−1^ due to a uniform increase in variation,*ψ*^−1^ for all genes.

In all cases, 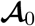 is 1000 × 15 sparse matrix such that column sparsity increases from 0% through 70% at a rate of 5% per column from the left most to the rightmost. The non-zero element of 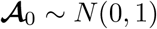. To induce hetero-geneity in source-specific variation relative to each other, 12 combination of (*ψ*_1_,*ψ*_2_), [(1, 10), (1, 7), (1, 5), (1, 3), (1, 2), (1, 1.5), (1.5, 1), (2, 1), … , (10, 1)] are chosen such that patients’ relative variance with respect to cell lines’,*ψ*_1_/*ψ*_2_ increases from 0.1, 0.14,0.2, through 10. For each pair of (*ψ*_1_,*ψ*_2_), we would like to vary shared variation in such a way that proportion of shared variation (*PSV*) increases gradually from 15% to 85%. One way to achieve such settings, is a) first define 10-point scale, *s* = 1, … , 10, then b) vary m by choosing an appropriate increasing function *f* on *s* such that *PSV*_*s*_ corresponding to *m*_*s*_ = *f* (*s,ψ*_1_, *ψ*_2_), goes from 15% to 85%. Note that since average total source-specific variation depends on *ψ*_1_ and *ψ*_2_, *f* also depends on them. To achieve this, we define the following form on 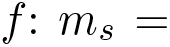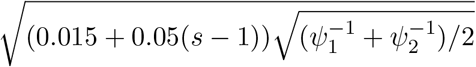. In all simulation settings to mimic Myeloma data we have set, *N*_*c*_ = 300, *N*_*p*_ = 200.

#### Impact of shared variation on iR_x_’s relative gain in accuracy

To measure the performance of the calibration models, we consider the predicted sum of squared errors (PSSE), defined as 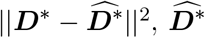 being the predicted response. PSSE measures the total unexplained error and is an estimate of the total MSPE across patients. Average PSSE’s of four methods are plotted in Figure S12 Section S11.3 of the Supplementary Materials. To have a comparable estimates across methods, which is scale independent, we focus on iR_*x*_’s relative gain (RG) in accuracy with respect to a given method, say *M*, defined as, 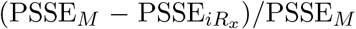. To summarize performance of iR_*x*_ we present 10 line-plots showing iR_*x*_’s relative gains corresponding to three methods, each for a fixed *PSV* against patients’ relative variance (*ψ*_1_/*ψ*_2_) in Figure 2.

**Fig 2:**
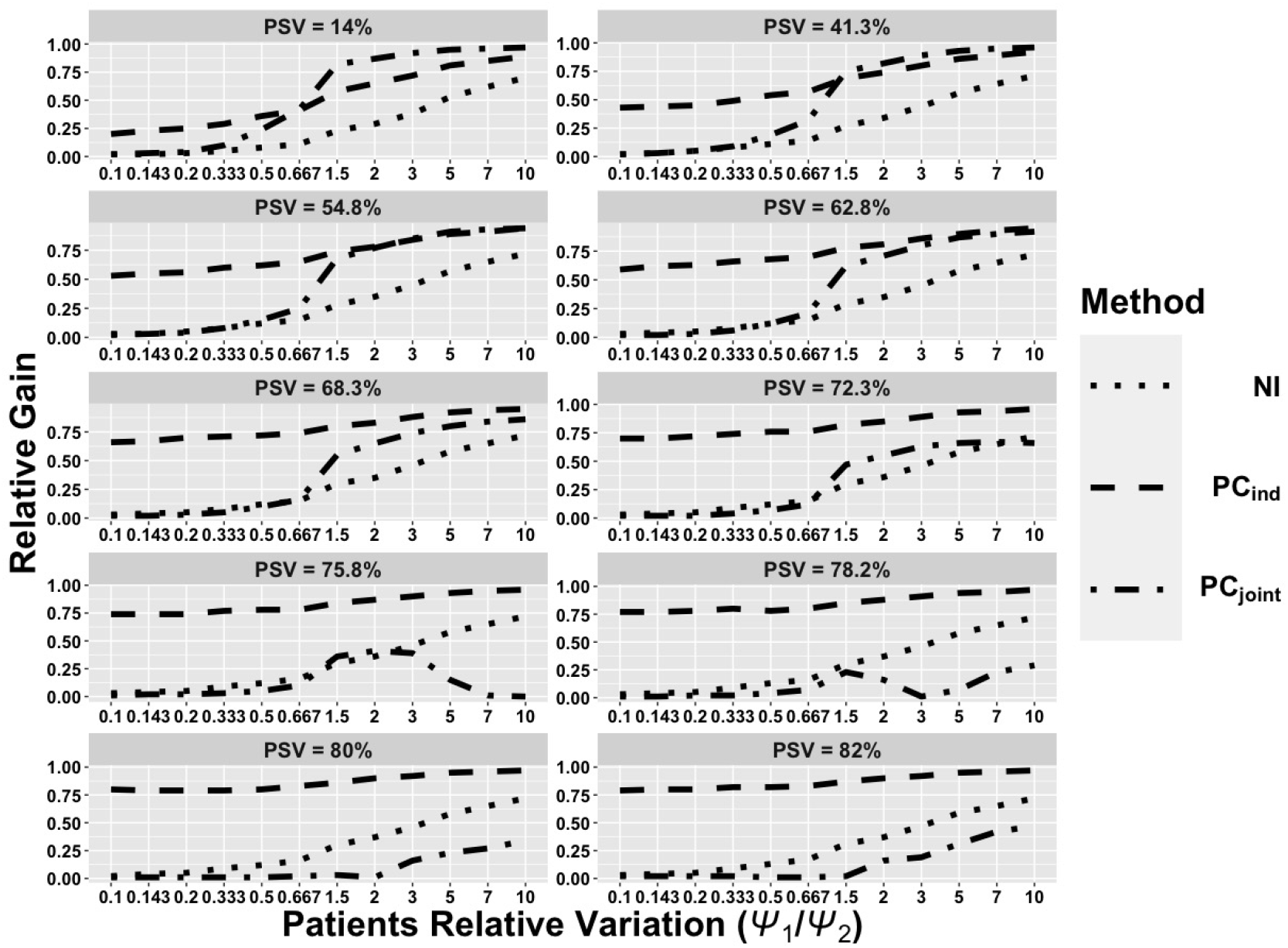
Line-plots for iR_*x*_’s relative gain(RG) with respect to three methods: NI, `PC`_ind_ and `PC`_joint_ (see text for explanation). Each of 10 sub-plots corresponds to a fixed *PSV*, that varies from 14% through 82% across the panels. In each sub-plot, the y-axis plots the RG against patients’ relative variance (*ψ*_1_/*ψ*_2_) in the x-axis.

Broadly, under the structure of some shared variation, RG of iR_*x*_ is always positive relative to all three methods and thus outperforms all competing methods considered. For cases when *ψ*_1_/*ψ*_2_ < 1 iR_*x*_’s RG remains low w.r.t both NI and PC regression methods while it increases as *ψ*_1_/*ψ*_2_ increases beyond 1 for almost all cases except for a couple of them where RG remains positive by a low margin for a few situations of (*ψ*_1_, *ψ*_2_) w.r.t `PC`_joint_. It is worthwhile to note that when *PSV* is low, NI model performs relatively better than PC-based methods as low shared variation approximates Naive case better. Though `PC`_ind_ consistently performs worse than other methods, `PC`_joint_ outperforms NI method for large *PSV*’s. It is noteworthy from Figure S12 (Section S11.3 of the Supplementary Materials) that for low *PSV*’s both PC regression methods perform considerably worse than NI and iR_*x*_ methods in terms of non-overlapping CI’s.

#### 4.1.2. Scenario II: Unknown structure of shared variation

Unlike earlier cases, here we assume the true models generating the values (gene expression) for **C** and **P** are not necessarily committed to a known structure of shared variation. We have followed two different approaches to simulate from NI model. In case a), we allow **C** and **P** to be generated from two separate processes. To achieve this, 50 replicates of 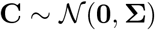, Σ ~ 50 * *IW* (Σ_*c*_, 50+*G*+1) whereas corresponding 50 replicates of 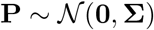, Σ ~ 50 * *IW* (**Σ**_*p*_, 50+ *G* + 1), where *IW* (.): Inverse Wishart distribution, 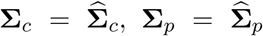, as estimated from the myeloma study. Note that these two *IW* have different mean matrix parameters. For simplicity,we say **C** ~ Scale Mixture of Normal or 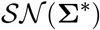 if 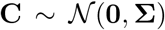, Σ ~ 50 * *IW* (**Σ**^*^, 50 + *G* + 1). In case (b), we assume both 50 replicates of **C** and 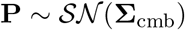, where 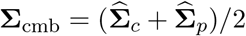. Although in case (b) generating process has the same mean matrix, there is good probability generated **Σ**’s are quite different from one another, and this variability spread across both **C** and **P**. In both cases, ***D*** is sampled as in Scenario **I** whereas ***ID***^*^ is simulated from *p*(***D*|C** = **P**). Boxplots of PSSE’s for all methods are given in the Figure 3.

**Fig 3:**
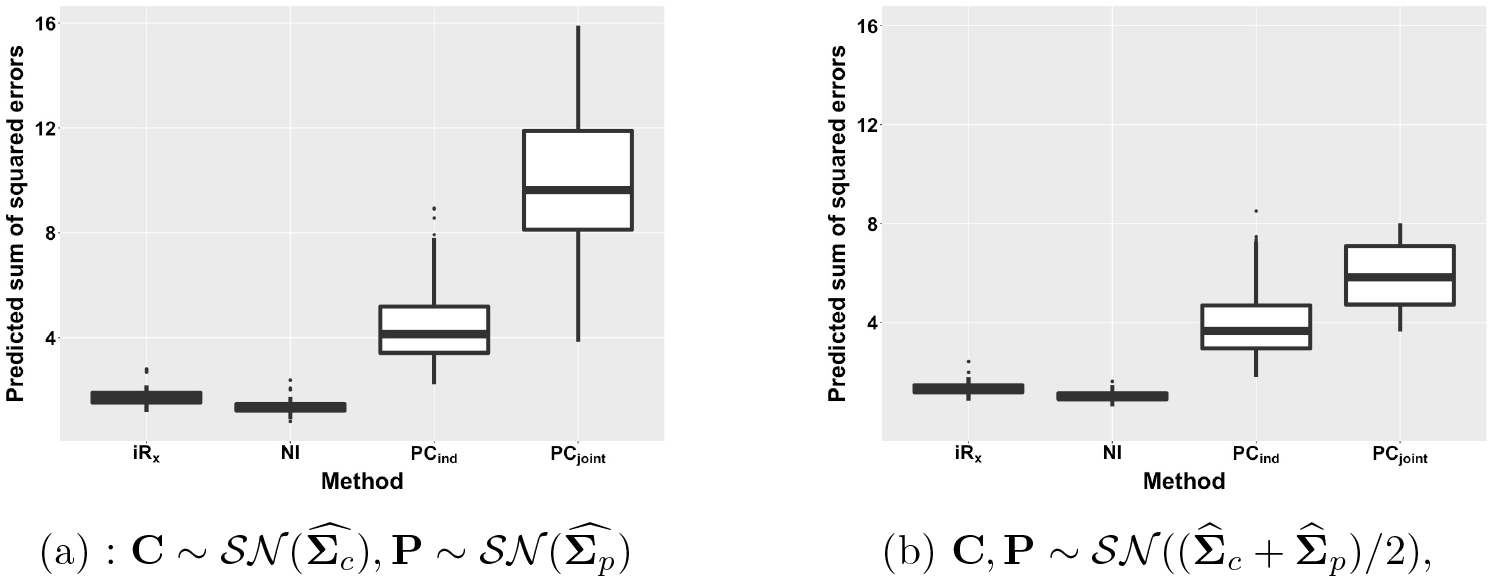
Prediction errors (PSSEs) of iR_*x*_,NI, `PC`_ind_, and `PC`_joint_ when the data generating model is mis-specified as NI: we have simulated from two types of NI models. Panel (a): 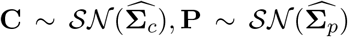; Panel(b): 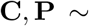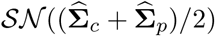.

The plot shows that iR_*x*_ is more robust compared to PC regression methods even when the true model is mis-specified as NI model. Since in this case, iR_*x*_ produces slightly biased estimates, as a result NI performs marginally better than iR_*x*_ on an average in terms of PSSE [case (a): 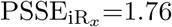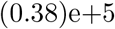, PSSE_NI_ = 1.37 (0.29) e+5, case (b): 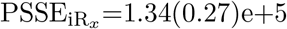, PSSE_NI_ = 1.02 (0.21)e+5]. However, on both the occasions, iR_*x*_ scores produce significant relative gains w.r.t PC regression methods [case(a): 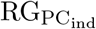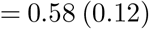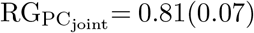, case(b): 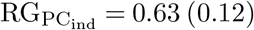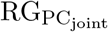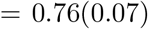 whereas both NI and iR_*x*_ scores, give forth more stable estimates than that of PC regression methods. Case(b) produces overall lower errors as the data were simulated from the same Inverse Wishart allowing them to have slightly more shared structure.

#### 4.1.3. Estimation accuracy of K, and stability of clusters

To measure the accuracy in estimation of *K*, and the stability of clusters induced, *K* and the clustering with cluster indices are recorded for each of 50 replicates under different simulation settings of (*ψ*_1_, *ψ*_2_, *PSV*) combinations as in scenario I. Given a simulation setting, to understand how stable the clustering is across replicates, we use average rand index (RI). Given a pair of clusterings of the same samples, say 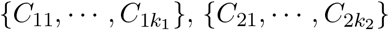,

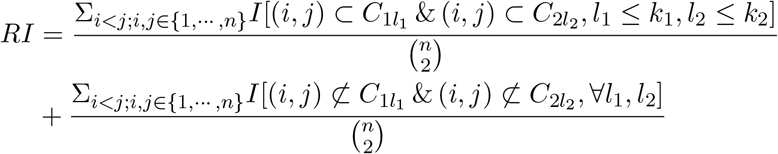

where *C*_*ij*_ is *j*^*th*^ cluster in *i*^*th*^ clustering. Thus, *RI* measures the proportion of concordant pairs, where a concordant pair (*i, j*) from samples is such that either 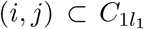 and 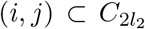 for some *l*_1_ and *l*_2_ or (*i, j*) does not belong to any cluster of either clustering. If two clusterings produce similar clusters, RI should be close to 1. *RI* = 1 for two identical clusterings. Once *RI* is computed for each pair of replicates, they are averaged over 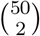 pairs. Both averaged *K* (with 95% CI), and averaged *RI* are reported in Table S4 and in Table S5 of the Supplementary Materials, Section S11.2 for each of different simulation settings as in Simulation Study I, Scenario I.

We note that *K* is estimated with quite high precision except for few cases corresponding to lowest proportion of shared variation (*PSV* ≈ 15%), which is not surprising as it usually leads to underestimation of *K* due to low signal. For all other cases (*P SV* > 35%), 95% CI always includes the true *K* allowing for continuity correction at boundary due to discrete nature of *K*^1^.For example, (*ψ*_1_ = 1, *ψ*_2_ = 3, *PSV* = 42%), *K*_est_ = 15(14, 16.25); (*ψ*_1_ = 3, *ψ*_2_ = 1, *P SV* = 41%), *K*_*est*_ = 15(15, 17.25); while (*ψ*_1_ = 1, *ψ*_2_ = 3, *P SV* = 15%), *K*_est_ = 10(8, 13); (*ψ*_1_ = 3, *ψ*_2_ = 1, *P SV* = 15%), *K*_est_ = 10.5(8, 13.5); We also point out that precision is quite good as average length of 95% CI across listed scenario is 2.84^2^. Next, to get a sense on how stable the clustering is across replicates, we observe a similar pattern as before. For case when *P SV* > 35%, average *RI* > 87%, whereas for cases when *P SV* ≈ 15%, *RI* remains relative smaller, but ≥ 80%. Thus, in presence of moderate to high shared variation, the iR_*x*_ methodology produces highly precise *K*, as well as, highly stable clusters.

### 4.2. Simulation Study II: Data-based simulation

Unlike Simulation Study I, with an aim to generate expression data, we mimic shared variation as estimated in Myeloma study. Thus, only deviation from the earlier settings is that we work with 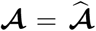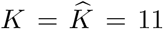, where both 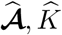 have been estimated from Myeloma study. Like before, we again consider 12 cases of (*ψ*_1_, *ψ*_2_) for the fixed amount of shared variation. For the first six cases,(*ψ*_1_ = 1, *ψ*_2_ ∈ {10, 7, 5, 3, 2, 1.5}) patients’ relative variance w.r.t cell lines’ 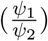 increases slowly from 0.1 through 0.67 whereas for next six cases, (*ψ*_1_ ∈ {1.5, 2, 3, 5, 7, 10}, *ψ*_2_ = 1) patients’ relative variance w.r.t cell lines’ increases relatively faster from 1.5 through 10. Since our objective is to compare the performance iR_*x*_ w.r.t NI, and to validate the claim on accuracy in prediction, we again consider PSSE’s and RG w.r.t NI as before.

Analogous to the previous section, we observed that iR_*x*_ always outperforms NI in prediction accuracy with respect to PSSE: as the patient’s relative variance increases, the RG increases monotonically. In the first six cases, the growth in accuracy remains slow (RG increases on an average from 3% through 14.9%); whereas, in last six cases, the relative gain grows very fast (from 28.2% through 71.8%). We further note that PSSEs fall for both iR_*x*_ as well as NI with the increase in *ψ*_1_/*ψ*_2_, estimates become more stable [first six cases: from (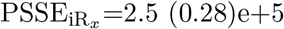, PSSE_NI_=2.58 (0.29)e+5) through 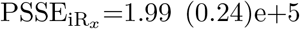, PSSE_NI_=2.34 (0.28)e+5)], last six cases: from [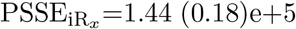, PSSE_NI_=2.0 (0.25) e+5] through 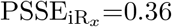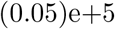, PSSE_NI_ =1.27 (0.18)e+5]. Both the phenomena can be attributed to the fact that data show very little evidence of significant improvement on incorporating patient-specific variations, while the patient-specific variation is extremely low. Otherwise, the consistent gain in accuracy validates the claim made in Section 2.3.

Next, we assess the performance of the iR_*x*_ model in real data-like setting where, unlike the previous case, all parameters are the estimates from the multiple myeloma study, including idiosyncratic source-specific variations and exact design parameters, such as *N*_*c*_ = 280, *N*_*p*_ = 188. Averaged 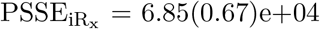, PSSE_NI_ = 8.77(0.94)e+04. Thus, we note that the relative gain in accuracy of iR_*x*_, with respect to NI, is 22%. Thus, the study once again confirms that iR_*x*_ outperforms NI, reestablishing the claim given in Section 2.3.

In the context of factor analysis, a positive lower triangular matrix constraint is usually imposed on the loading matrix to prevent the standard identifiability issue (Geweke and Zhou, 1996; Lopes and West, 2004). As suggested in De Vito et al. (2018), we also applied Orthogonal Procrustes (OP) algorithm (Aßmann, Boysen-Hogrefe and Pape, 2016) based on the post-MCMC processing approach to ensure identifiability of 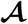. Under the unconstrained setting, the samples obtained from the posterior distribution of 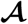 are simply an orthogonally rotated variant of the unique estimator of the parameter 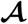. Throughout the paper, we carry forward our analysis using the posterior mean of 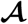 and the uniqueness of 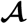 is ensured through multi-modality test (Gelman Rubin statistics, Gelman and Rubin (1992); Brooks and Gelman (1998)), convergence test (Geweke test, Geweke (1991)) along with an implementation of rotation invariant OP algorithm. More details about the identifiability of 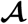 in simulation and real data settings are deferred to Section S5 of the supplemental file.

## 5. Clinical validation using existing trial data

We perform an *in silico* validation of our iR_*x*_ model based on retrospective clinical trial data. To validate our approach, we identified existing clinical trial datasets that had collected gene expression measurements before drug treatment and had subsequently measured a clinical drug response phenotype. Our primary goal here is to compare the calibrated iR_*x*_ estimator to the <u>true</u> drug responses from the patients (not used for fitting). This is to assess whether our integrated models derived from patients and cell-lines can explain a significant proportion of the actual drug responses from patients. For this purpose, we consider patients’ transcriptomic (gene expression) measurements from two clinical trials, namely administering bortezomib on myeloma patients (Mulligan et al., 2007) and docetaxel on breast cancer patients (Chang et al., 2003). To train the model, we use cell lines’ gene expression and pharmacological drug response data from Cancer Genome Project (CGP, Garnett et al. (2012)), whereas patients’ gene expression data are derived from the Gene Expression Omnibus (GEO) database (Clough and Barrett, 2016), in which data were contributed by the authors of the corresponding clinical trials. The GEO also contains the drug response data, along with the tissue microarray data. We preprocess the datasets to obtain drug response data across cell lines and gene expression for the same set of genes for both cell lines and patients following Geeleher, Cox and Huang (2014). Details of data processing are described in the Supplementary Materials (see Section S6 and the provided R-codes). For each clinical trial, we fit our iR_*x*_ model and calibrate patients’ drug response. To assess the prediction accuracy of our model, we compare scores obtained from all four models, namely, iR_*x*_ NI, `PC`_ind_ and `PC`_joint_. We then discuss utilities of our clustering approach by interpreting mixed clusters in terms of drug response phenotypes, finding key genomic drivers of the shared variation, and exploring their association with potentially relevant functional pathways.

### 5.1. Bortezomib study in Multiple Myeloma Patients

Therapeutic choices for myeloma have become increasingly complex as the optimal use of new agents remains unclear, either alone or in combination with standard therapy, and standard clinical prognostic factors are unable to reliably identify the highest risk patients (Mulligan et al., 2007). Under these circumstances, providing a precise genomic classifier will help clinicians understand and predict the outcome for a specific therapy. Bortezomib is the first proteasome inhibitor approved by the US Food and Drug Administration (FDA) for the treatment of multiple myeloma and relapsed/refractory multiple myeloma (Kane et al., 2003). We apply our methodology to a large study of relapsed multiple myeloma patients, who enrolled in phase II and III clinical trials of bortezomib and consented for pretreatment gene expression profiling using tissue microarrays. For each patient, authors initially defined clinical response into five categories (i.e., Complete Response (CR), Partial Response (PR), Minimal Response (MR), No Change (NC), and Progressive Disease (PD)) based on the degree of effectiveness after no more than eight cycles of bortezomib. Later, they further grouped MR, PR, and CR as responders (n=85), and PD and NC as non-responders (n=84). For the training model, 280 samples of drug perturbation data of bortezomib on cell line model systems along with their gene expression data are independently collected from the GDSC platform, as used in Geeleher, Cox and Huang (2014)’s study. We chose the top 1000 genes that show high (marginal) correlation with drug responses from our cell-lines for the analysis.

### Improved therapeutic response calibration

First, we note that responders are expected to register lower iR_*x*_ scores (congruent to IC_50_ values) than non-responders, as a lesser amount of a sensitive drug is needed to achieve the same effect as a resistant drug. The violin plots in Figure 4(a) show that standardized iR_*x*_ scores are shifted downward for responder as desired, unlike NI scores, which show little difference in their distribution across two response groups. Our proposed model iR_*x*_ provides higher prediction accuracy compared to other three models in terms of p-values from one-sided t-tests between responder vs non-responder, (iR_*x*_=0.0245) versus (NI=0.8),(`PC`_ind_=1), and (`PC`_joint_=0.97) respectively, with iR_*x*_ giving forth the only significantly different group means. We also use another measure, the area under the receiver operating characteristic curve (AUC), to compare the scores between the groups. A receiver operating characteristic (ROC) curve is a plot of the true positive rate (sensitivity) as a function of the false positive rate (1-specificity) for different cutoff points of a predictor. The higher the AUC, the better the prediction rule in distinguishing the two groups. iR_*x*_ outperforms all three other models in terms of AUC (0.59 versus 0.46,0.39 and 0.41 respectively). One of the reasons PC-based methods perform worse than iR_*x*_ is that unlike PC-based methods iR_*x*_’s prediction mechanism takes into account source specific variations, which is quite large in this case, and causing other purely mean-based prediction methods to fail. To measure how iR_*x*_ does as a dichotomized rule that tells responders apart from non-responders for a clinician, we use the c*σ* rule (see Section 2.3) ignoring samples predicted as “No change.” We evaluated the value of *c* for both the clinical trials under consideration and recommend a conservative value of *c* = 0.5. Note, our method works well even for smaller values and does not impact the inferences significantly (see Section S7 of the Supplementary Materials). For measuring the performance, we compute the enrichment probability (EP), which is the posterior probability that the proportion of responders restricted to the class of predicted responders is bigger than that of non-responders, assuming priors on the proportions as Beta(1,1)’s (see Section S8 of the Supplementary Materials for more details). The closer the EP is to 1, the stronger the evidence that the classifier’s predicted categories for responders and non-responders are enriched by the true ones. The EP score for the *cσ* classifier is 0.99, where c=0.5. Approximately 60% of predicted responders are true responders and 62.5% predicted non-responders are true non-responders, which is encouraging.

**Fig 4:**
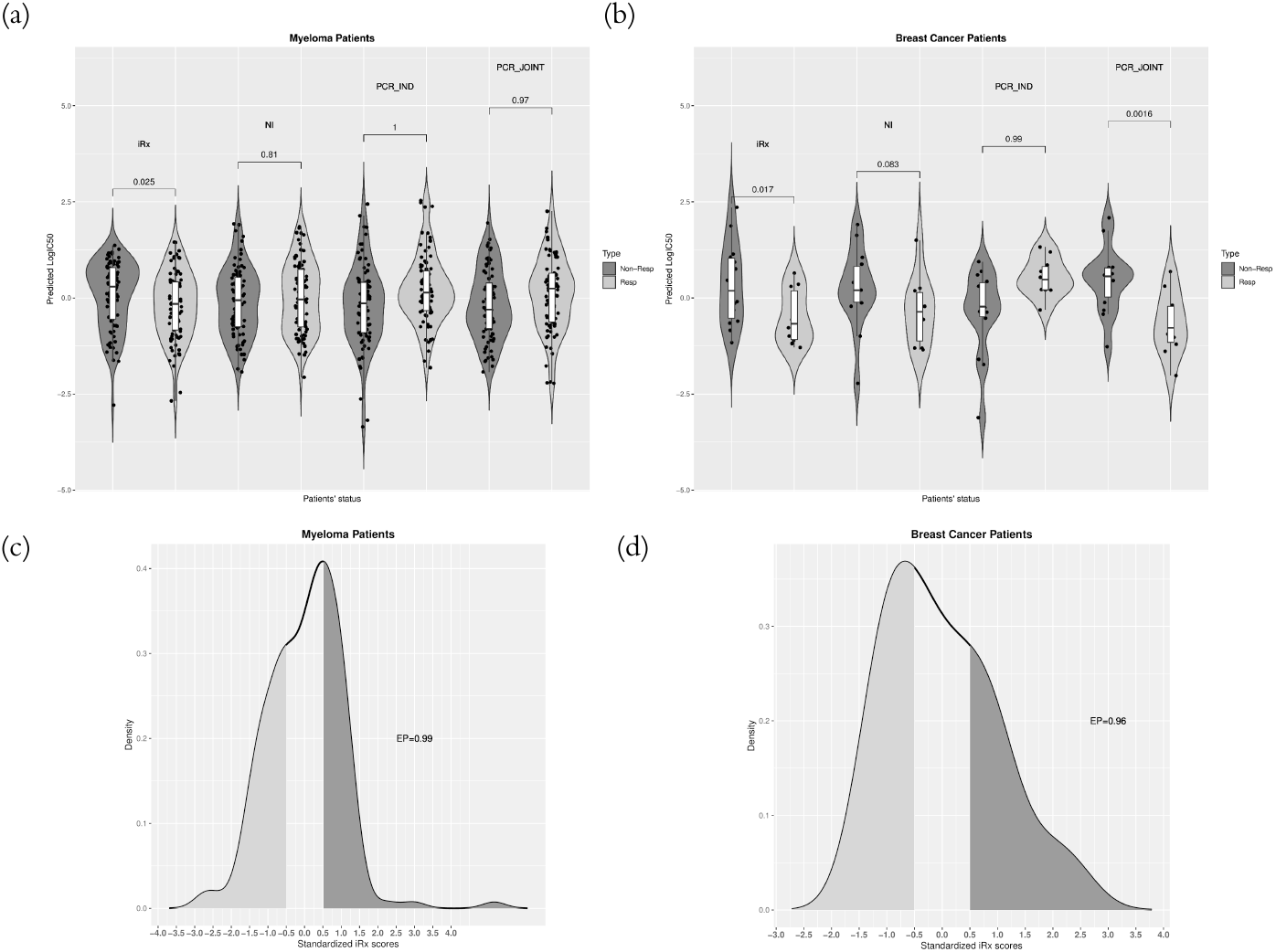
The performance of iR_*x*_ is compared with three models, namely, NI, `PC`_ind_ and `PC`joint for Panel (a) Multiple Myeloma study and Panel (b) Breast cancer study. In both (a) and (b), each pair of violin plots of non-responders (dark grey) and responders (light grey) represents standardized continuous scores of iR_*x*_, NI, `PC`_ind_ and `PC`_joint_ from the left to the right respectively. In the bottom panels, the dichotomized iR_*x*_ *cσ* classifier(c=0.5) is demonstrated on the density plot of standardized iR_*x*_ for Panel (c) Multiple Myeloma study and Panel (d) Breast cancer study. The shaded region represents the predicted category along with the enrichment probability (EP).

### Cell lines as patient avatars

We have found 11 clusters, denoted by {*C*_1_,*C*_2_, … , *C*_11_}, based on the 11 estimated factors from all the samples of patients and cell lines: among them, 8 clusters include both cell lines and patients (*C*1 − *C*4, *C*6, *C*8 − *C*10). Based on the distribution of predicted iR_*x*_ scores across the 169 patients, we further have classified the patients into three groups: *neutral* for patients who have the predicted score in the symmetric region around 0, accounting for approximately 30% of the 169 scores; *responder* for patients who have lower values than *neutral*, and *non-responder* for patients who have higher values than *neutral*. Figure 5 displays barplots of the cluster sizes and shows the proportions of cell lines and responders/non-responders/neutral patients within a cluster. There are three homogeneous clusters: *C*5 has only cell lines, whereas *C*7 and *C*11 include only responder and non-responder patients, respectively. The cell lines in mixed clusters can be determined as avatars representative of the patients with respect to potential drug response effectiveness. We are particularly interested in the avatars of the mixed clusters having large predicted responders. To better represent them pictorially we present a chord diagram (Gu et al., 2014) in Figure 5 (b) that displays relations between avatar cell lines and patients. The gray and purple sectors around the circle represent cell lines and patients, respectively. The purple sector is split further into the drug response groups, namely neutral, responder, and non-responder. The connections in the same color represent links between cell lines and patient groups in the same cluster. The avatars are highlighted in Figure 5 (b).

**Fig 5:**
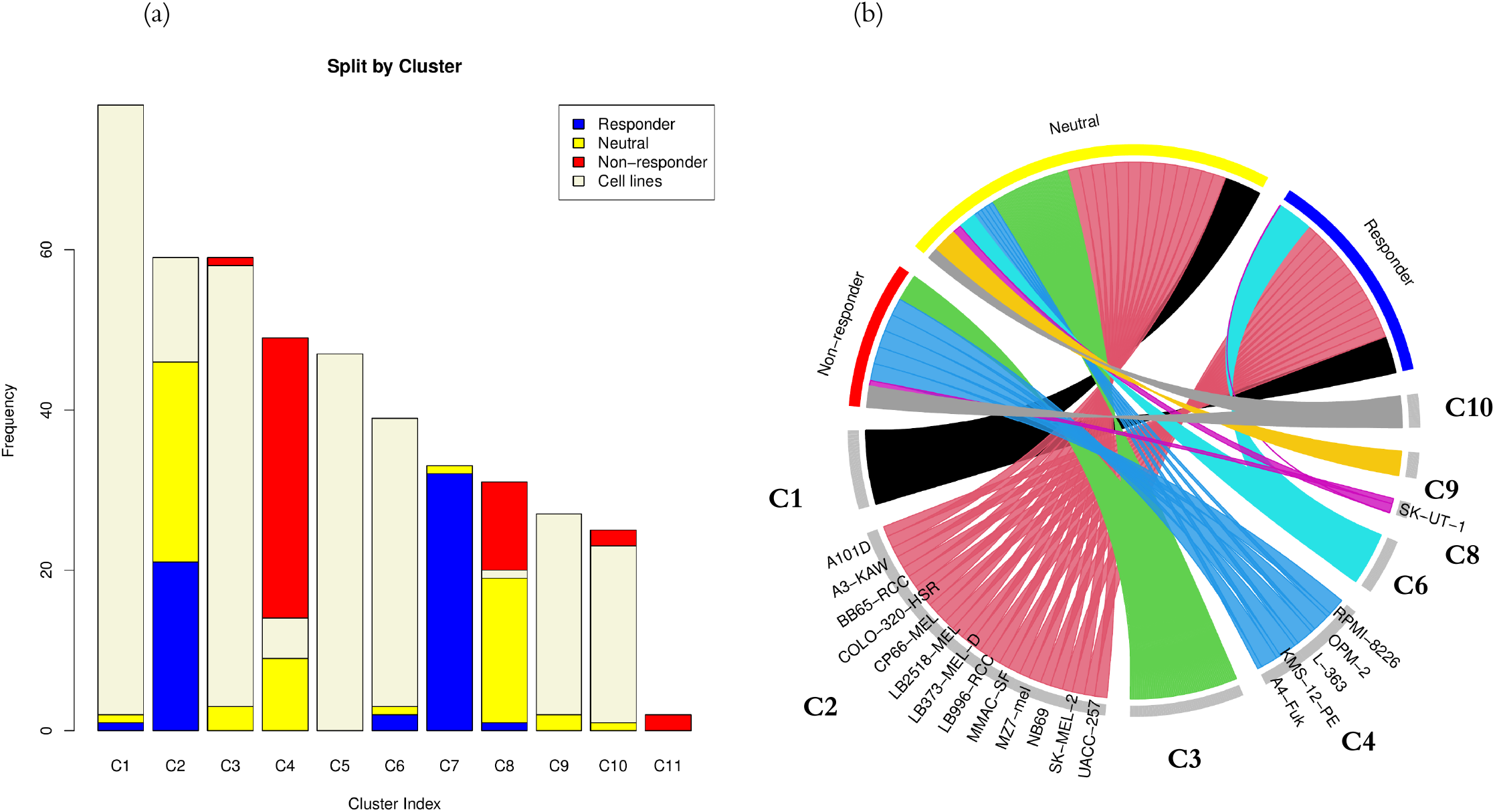
Clusters of samples in the multiple myeloma study, and cell lines as avatars for patients: (a) gives distribution of cell lines and three subtypes of multiple myeloma patients (responder, non-responder, and neutral) across clusters, arranged in order of cluster size, whereas (b) depicts colored links (chords) between cell lines (gray sectors) and patients’ samples within clusters. The total width of the chords originating from a cell line, which is proportional to number of patients in the cluster, defines link strengths. Cell lines with high total chord width are considered important avatars as highlighted in *C*2, *C*4 and *C*8.

Interestingly, the predictive scores in the responder patient group are mostly represented by cell lines in *C*2. Among the avatars of *C*2, *BB65-RCC, A101D* and *A3-KAW* are particularly remarkable as they are extremely sensitive to bortezomib having a low relative sensitivity Z-score ^3^ (< −1.5), as obtained from the portal of Genomics of Drug Sensitivity in Cancer (GDSC, Yang et al. (2012), https://www.cancerrxgene.org/^4^). Though the avatars may represent various tissue-types, there are multiple established renal cell carcinoma cell lines (*BB65-RCC, LB996-RCC*) and melanoma cell lines(C*P66-MEL, LB373-MEL-D, MZ7-MEL, SK-MEL-2*) among them. To this end, previous studies indicate that bortezomib can be very effective in sensitizing certain renal cell carcinomas (Brooks et al., 2010) while it can kill melanoma cancer cells when synergized with other drugs (Yerlikaya et al., 2010; Reuland et al., 2012). Similarly, *C*1 and *C*6 both have a small section of responders compared to the larger set of cell lines. As a result, though there may not be clear candidate(s) for avatar(s), *BE-13,DEL,DB* for *C*1 and *DSH1* for *C*6 are found to be quite sensitive to bortezomib, thus posing to be avatar candidates. This shows that avatars have the potential as good pre-clinical models for the respective patient subgroups. For the cluster of *C*8 we see a mixture of both responders, non-responders, which is represented by a sole avatar, namely *SK-UT-1*: an established uterus cell line. It gets a Z-score of −0.266 for bortezomib, which is considered neither sensitive nor resistant, thereby validating the clustering. The scores in the non-responder group are predicted by cell lines in *C*4 and *C*10. It turns out the avatars are found in *C*4 and all of them except *A4-Fuk*, namely *KMS-12-PE,RPMI-8226,OPM-2,L-363* are established myeloma cell lines. They all have a Z-score > −0.89 except for *KMS-12-PE*, for which a Z-score is not registered with the GDSC portal. Thus, again we note that clustering, which is carried out independent of drug response study, seems to bring together patients and cell lines such that patients’ predicted responses are similar to the observed responses of the cell lines.

### Driver genes and pathways

Driver genes are selected based on gene-factor relationship from the standardized factor loadings, *L*_*m,h*_ (as in Section 3.5). Figure 6(a) displays a circular heatmap of the significant factor loadings *L*_*m,h*_ for *K* = 11 latent factors, labeled as F1 through F11. For the most sparsely loaded factor F11, the five most significant genes (namely, *SLC39A7, GGCX, KDELR3, PLEC, and TM9SF1*) can be easily identified in the reddish-yellow strip at the one o’clock position on the outermost layer, which appear together due to clustering.

**Fig 6:**
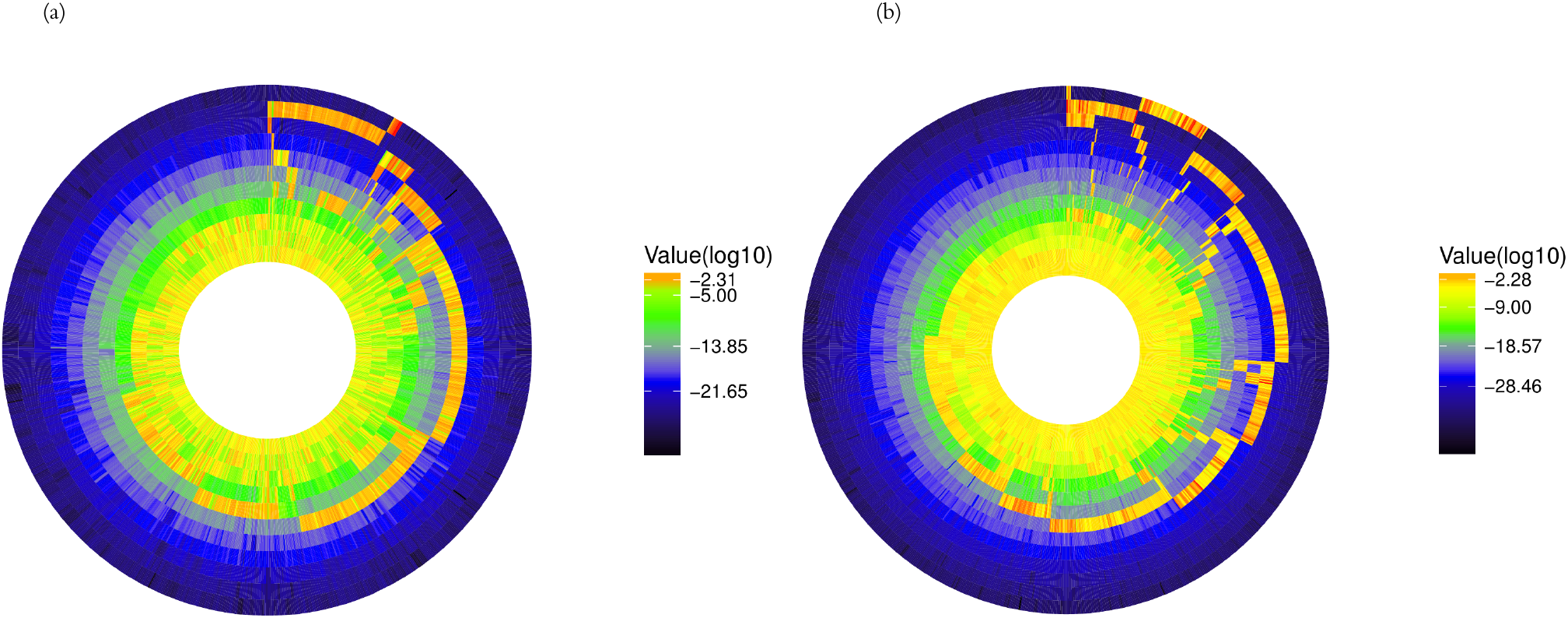
Circular heatmap of standardized loadings (*L*_*mh*_): The circular plots represent genes’ loadings’ contribution to latent factors for (a) multiple myeloma with 11 latent factors, and (b) breast cancer with 14 latent factors. In both cases, the innermost circle represents the first factor (F1) and then F2, F3, and so on outwards. The relative sparsity increases as we move toward the outermost circle (Note that black/blue regions represent extremely low values, whereas yellow/red regions represent high values)

For each of the selected gene sets, a pathway analysis is carried out using the Ingenuity Pathway Analysis tool (Krämer et al., 2013). The analysis compares the selected gene set with reference gene sets corresponding to pathways (Ingenuity Knowledge Database (genes only) is used for reference sets) by conducting the Fisher’s exact test. The tool outputs pathways in order of adjusted p-values, which make them comparable. To understand which pathways are potentially most relevant given a factor, a wordcloud is constructed with keywords selected from “significant” pathway names. A wordcloud highlights the keywords based on its relative frequency of occurrences in a text. Figure S9(a) (See Section S9 of Supplementary Materials) shows such a wordcloud generated from keywords from pathway names corresponding to a p-value cut-off of 20% ^5^, where the sizes of the texts correspond to the frequencies of keywords, and M1,M3 etc represent the clusters of pathways corresponding to the underlying factors F1,F3 etc. Some of the highlighted keywords, such as *MAPK* for M1, *Endocytosis* for M4, may point to the potentially relevant pathways. For example, in this case, they may indicate Mitogen-activated protein kinase (MAPK) signaling pathways and the endocytic pathways, respectively. A signaling pathway describes a group of molecules in a cell that works together to control one or more cell functions, such as cell division or cell death. After the first molecule in a pathway receives a signal, it activates another molecule whereas endocytic pathways initiate a cellular process, called *Endocytosis*, in which substances are brought into the cell. Martin-Antonio et al. (2015) showed that NKG2D (a receptor activating natural killer cells (NK)) expression, in multiple myeloma (MM) cells after cord blood-derived natural killer cell (CB-NK) treatment correlates with lower MM progression, and NKG2D and NKP30 contribute more to the cytotoxicity of MM cells. In this regard NKG2D and NKP30 are transferred into MM cells via lipid trafficking and the endocytic pathways, thereby helping CB-NK cytotoxicity. A full list of all significant pathways are provided in Table S2 in Section S9 of the Supplementary Materials.

### 5.2. Docetaxel study in breast cancer patients

Docetaxel is a new class of cytotoxic agent approved for breast cancer (Crown, 2001). We consider 24 breast cancer tumor biopsies through a clinical trial, which measure the response of patients to docetaxel in a neoadjuvant setting (Chang et al., 2003). In this study, percentage reduction of tumor size, measured after four cycles of docetaxel, is used to measure effectiveness of the drug. The authors designate a patient as a responder (n=10) if the tumor shrinks to <= 25% at the end of the study period, otherwise as a non-responder (n=14). Because of low patient-size, a predictive model for drug response cannot be trained solely based on part of the samples. Such scenarios are very common in clinical trials and lay the foundation for the need of our integrative modeling framework. For training Cell line-patient model, 482 samples of drug perturbation on cell line model systems along with their gene expression data are independently collected from GDSC platform.

### Improved therapeutic response calibration

Our proposed calibration index, iR_*x*_, captures more variation in drug response phenotype compared to NI and `PC`_ind_, while performs at par with `PC`_joint_ (Figure 4(b)). The iR_*x*_ score outperforms NI and `PC`_ind_, both in terms of p-values (iR_*x*_= 0.017, NI= 0.08, `PC`_ind_=0.99) and AUC (iR_*x*_= 0.75, NI=0.71, `PC`_ind_= 0.23), while it performs at par with `PC`_joint_(p-value= 0.016, AUC=0.83). Though `PC`_joint_ seems to be marginally better, iR_*x*_, `PC`_joint_ and NI may be deemed similar due to low power (*n* = 24). Since unlike `PC`_joint_, `PC`_ind_ uses one SVD loading matrix based on C for estimation of regression parameters while another SVD loading matrix (say 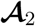) based on **P** for prediction, the inferior performance can be attributed to poor estimation of 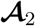 due to low sample size of pa-tients as true 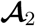, which can be treated as an empirical factor loading matrix (see West (2002)), is closer to common 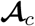 based on [**C, P**] as is the case with `PC`_joint_. In this case, the *cσ* iR_*x*_(c=0.5) classifier yields an EP score of 0.96. Around 60% predicted responders are true responders, whereas around 85.7% predicted non-responders are true non-responders.

### Cell lines as patient avatars

The iR_*x*_ model has identified 14 latent factors and thus produced 14 clusters, namely *C*1 through *C*14. Clustering results are displayed in Figure S10 in Section S10 of the Supplementary Materials. We note that patients are located in *C*1 − *C*5 and *C*14; among which *C*14 is a homogeneous cluster of patients, predicted to be non-responders only. The mixed clusters *C*1 − *C*3 were predicted to have mostly responders, while *C*5 consists of mostly predicted non-responders. As evident in Figure S5 (b), the chord diagram does not highlight avatar cell lines due to small size of patients with respect to the size of cell lines. Nevertheless, from clinical standpoint it is worthwhile to investigate *C*1 − *C*3 for potential candidates for avatars as they consist of predicted responders only. In *C*1, bortezomib is found potent against *TE-8, CAL-62, GAMG, HT-1080* and *Hs-578-T* (having a relative sensitivity Z-score ~ −1.8) out of which only *Hs-578-T* came from breast cancer, namely invasive ductal carcinoma. 4 cell lines, namely *SF539,FTC-133,D-566MG, BB49-HNC* with non-breast cancer origins are found to be sensitive to bortezomib in *C*2. In particular, the drug is extremely potent against *FTC-133* (a thyroid cancer cell line) and *D-566MG*, (a Glioblastoma cancer cell lines), each having a Z-score < −2. *C*2 and *C*3 contain 5.6% and 14.5% breast cancer cell lines respectively and almost all of them show no change (or neutral) to docetaxel.

### Driver genes and pathways

Figure 6(b) provides a circular heat-map of the significant factor loadings *L*_*m,h*_ for K=14 factors to show gene-factor relationship. Similar to myeloma case, we perform an IPA analysis and construct a wordcloud for breast cancer study as given in Figure S9(b), where D1,D2 etc represent the clusters of pathways corresponding to the underlying factors F1,F2 etc. The plot indicates top keywords, such as, *lymphocytes* for D5,D12; *macrophages* for D13; etc. The keyword, macrophages, might be related to inhibition of Stat6 pathway. Binnemars-Postma et al. (2017) argued increasing evidence has demonstrated a relationship between a high degree of macrophage infiltration and poor prognosis in human breast cancer and other malignancies, which suggests that macrophages play an important role in tumor progression and metastasis in cancer. He later points out that inhibition of the Stat6 pathway in tumor-associated macrophages (TAMs) is a vital therapeutic approach to attenuate tumor growth and metastasis by inhibiting TAM-induced protumorigenic and prometastatic activities. Table S3 in Section S9 of the Supplementary Materials lists significant pathways that may be associated with these factors.

## 6. Discussion and future directions

In this article, we propose an integrative Bayesian framework, called the iR_*x*_ model, that integrates both labeled and unlabeled data across patient, cell line models systems, and drug perturbation platforms by defining a joint probability model. The semi-supervised formulation allows computation of therapeutic response to new patients exploiting the genomic similarity between patient and cell lines model systems – to aid implementation of evidence-based personalized medicine strategies. Bayesian inference via explicit sub-models enjoys interpretable and computational properties; such as, a) latent factor model allow effective dimensional reduction, b) sparse linear Gaussian framework admits fast and closed-form computation and calibration iR_*x*_ distribution and scores. In addition, the unified Bayesian framework proposes fast computation for clustering both patient and cell line samples based on shared variation. Cell line-patient model (C-P model) is particularly useful to identify conserved biological mechanisms between genomic profiles with two different types of sources-specific variations.

We have established through both simulation studies and real data settings, namely a bortezomib with multiple myeloma study and the docetaxel on breast cancer study, that our proposed novel therapeutic index calibrates responses with higher precision than the standard alternatives. We have provided some theoretical support for this gain in accuracy through explicit modeling of shared variation. It can be further shown that iR_*x*_ scores are robust to the change in number of features in real data settings, which is not the case with usual ridge regression models. This is due to the fact that changes in the key factors happen at a slower rate with increasing number of features. In real data settings, we have clustered samples and defined avatar cell lines representative of patient subgroups that confirm existing knowledge and provide insights into novel oncogenic mechanisms. We have supplied all the data and code required to reproduce the results of this paper along with the Supplementary Materials.

There are several refinements and generalizations of our model that are possible. For example, the C-P model can be extended to any number of other pan-omic platforms, such as proteomics, metabolomics, and epigenomics with platform-specific precision matrices. The inclusion of more platforms may improve therapeutic indices, as a comprehensive set of molecular features is more likely to better capture key underlying oncogenic features, although it can potentially increase the computational burden. From a methodological perspective, in our iR_*x*_ model formulation, we assume the source-specific variations are consumed in the error terms (**Ψ**_•_). We do so for multiple reasons: first, our primary objective is to infer the shared latent factors (through 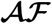) which informs our final prediction model. Second, this construction allows us to explicitly calibrate the iR_*x*_ distribution using closed form expressions (as shown in Section 2.3) and more importantly, quantify the gains in prediction accuracy (as shown in Proposition 1). This assessment is further (profitably) validated both in our simulation studies and real clinical data examples. Finally, this formulation is also computationally feasible since we are estimating one set of (shared) latent factors. In the context of factor modeling of expression data from high-throughput platforms, the current literature has some versatile formulations in terms of (explicitly) capturing source-specific variations such as those proposed by De Vito et al. (2019). However, it introduces an additional layer of complexity both in terms of model construction (through the incorporation of additional covariance terms) and ensuing computational burden (estimation of corresponding multiple latent factors). While no doubt interesting, we eschew this in our current work for the above reasons and leave this as a future extension.

Currently, the drug-cell line model is specified by a linear Gaussian framework, which can be extended to a non-linear framework, however this may not allow explicit closed-form computation of iR_*x*_ scores. While proposing the iR_*x*_ score, we have also considered a dichotomized c*σ* classifier to determine if a patient is a potential responder. *σ* is computed empirically based on the calibrated estimates. One might be interested in proposing a standardized score based on a *σ*, which is model dependent instead. To arrive at such a *σ*, one can consider 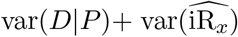, which might require further investigation.

Another interesting avenue to pursue would be where cell lines are allowed to belong to multiple clusters using soft clustering methods as opposed to the hard clustering methods we propose in this paper. For example, there has been recent work (Liu et al., 2020), which used a multi-scale network-based clustering approach to identify patient-specific clusters of cell-lines and evaluated the connectivity using spectral wavelet decomposition on graphs. Another possible extension of the *K*-means with additional parameter called “fuzzifier” that determines the level of cluster fuzziness, e.g. Fuzzy C-means algorithm (Bezdek, Ehrlich and Full, 1984) can be applied to the estimated factor matrix and the continuous type membership coefficients of cell lines can be used as weights for patient groups. The representation of avatar cell lines for multiple patient groups will increase utilities of our iR_*x*_ framework and can be easily incorporated as a post-iR_*x*_ model fitting procedure.

The core construction of our iR_*x*_ framework can be applied to a broad range of problems in data integration and application areas. From a methodological perspective, our semi-supervised models can be used and adapted to combine multiple sources of data, where-in some of the data sources have missing/unlabeled outcomes. This includes not only other biomedical settings (e.g. Boonstra, Mukherjee and Taylor (2013)) as well as electronic health records data (e.g. Gronsbell and Cai (2018)). From an applications perspective, in pharmacogenomics, diverse model systems of human cancer such as immortalized cancer cell lines, either cultured *in vitro* or grown as xenografts, and patient-derived xenografts (PDXs) or organoids are being extensively used for the development of cancer drugs as well as understanding both biological mechanisms and clinical responses. Specifically, PDXs are created by implanting tumor cells from patients in mice and have been proven to accurately reflect the patient’s tumor properties. Such co-clinical avatar studies allow for real-time adaptive therapeutic decisions by finding the best-matched PDX for a given patient based on molecular profiling data then these PDX samples are used to inform the patients’ treatment options (Byrne et al., 2017). While there has been significant research on conservation of genomic profiles between PDX and patient tumors, little work has been done for providing statistical integrative systems to inform patients’ drug response – which is precisely the gap our work aims to fill. Motivated by such co-clinical designs, in our recent studies (Bhattacharyya et al., 2020; Liu et al., 2020), we focused on accessing cell lines as representative models for patients to understand patients’ drug sensitivity. As a continuation of these efforts, we plan to apply iR_*x*_ to PDX data as training model system to calibrate patients’ therapeutic outcomes, as these data mature in the coming years.

In summary, drug re-purposing, or repositioning is an approach that can significantly reduce the time, cost, and risk of drug development, which involves the application of existing drugs (approved or under experimentation) for the treatment of previously unconsidered indications. Thus, given a set of (potentially large number of) drugs being considered for a particular treatment, our iR_*x*_ model can be applied to calibrate and refine therapeutic indices on patient populations – which can aid prioritization of candidates (or combinations) for personalized therapy regimens.

## Acknowledgements

We would like to thank the Editor, the Associate Editor and two anonymous referees for their constructive comments and references that led to significant improvements of the article.

## Supplementary Materials

Supplementary Materials to “A Bayesian Precision Medicine Framework for Calibrating Individualized Therapeutic Indices in Cancer” contains all additional mathematical derivations, proofs, additional results and figures as well as all supporting data and codes are attached separately.

## APPENDIX A: TABLE OF NOTATION

## Footnotes

1 since we are considering discrete intervals, median estimate can lie on the boundary. One way to avoid that is to use continuity correction at the boundary

2 includes continuity correction

3 Z-score is a logarithm of IC50 value standardized across all cell lines the drug is applied to. Z-score being close to −2 or less is indicative of extremely potent drugs while having > 2 values indicates a resistant drug. Any value in between them indicates the drug is neither sensitive nor resistant.

4 The website lists two types of sensitivity Z-score, namely GDSC1 and GDSC2, based on types of design and assay. GDSC2 represents a more modern method and thus, is used here. GDSC1 is used wherever GDSC2 data are not available.

5 A conservative choice of 20% is considered to guarantee large set of pathways representing all of the factors

## Notes

† Supported by National Institutes of Health Grants No. R21CA220299- 01A1 (to M.J.H. and V.B.), U54-CA224065, and 3P50CA070907- 20S1; Leukemia and Lymphoma Society Grant No. 7016-18; Cancer Prevention and Research Institute of Texas Grants No. RP180712, (to M.J.H.), R01-CA160736, R01-CA194391, R01CA244845-01A1 and P30-CA46592; National Science Foundation grant DMS 1922567; funds from the UM Rogel Cancer Center and the School of Public Health (to V.B.)

### Competing Interest Statement

The authors have declared no competing interest.

